# Dissecting the Heterogeneity and Tumor-Associated Dynamics of Human Liver Group I ILC via scRNA Sequencing Data

**DOI:** 10.1101/2024.10.24.619932

**Authors:** Yumo Zhang, Jitian He, Xue Li, Huaiyong Chen, Zhouxin Yang, Youwei Wang

## Abstract

Single-cell transcriptome analysis has made outstanding contributions to the identification of new cell lineages and the study of cancer immune microenvironment. Yet, the characterization of human liver ILC1s and their dynamic changes in the tumor microenvironment have not been thoroughly studied at this detailed level. Here, we performed an integrated analysis of mouse and human liver immune cells to identify human liver ILC1 based on identified mouse liver ILC1 and to verify its functional similarity. Additionally, our findings highlighted the different expression patterns of the EOMES gene in human versus mouse group I ILCs, suggesting its reduced regulatory significance in human liver NK cells and ILC1 compared to murine models. A unique subset of *CD127*^hi^ intermediate ILCs (intILCs) was identified, exhibiting traits of both human liver NK cells and ILC1. Single-cell sequencing data analysis and TCGA were utilized to characterize the distinct alterations in the genes and functions of NK cells, ILC1, and intILCs in human hepatocellular carcinoma. It was found that the dynamic changes of ILC1 and intILCs, along with some of their subpopulations, may be key factors in tumor progression. This study provided new insights into the identification of ILC1 in human liver and the immunologic changes and mechanism of group I ILCs in the tumor microenvironment, and these findings may be applicable to improving the diagnosis and treatment of hepatocellular carcinoma.

**Graphical Abstract:** 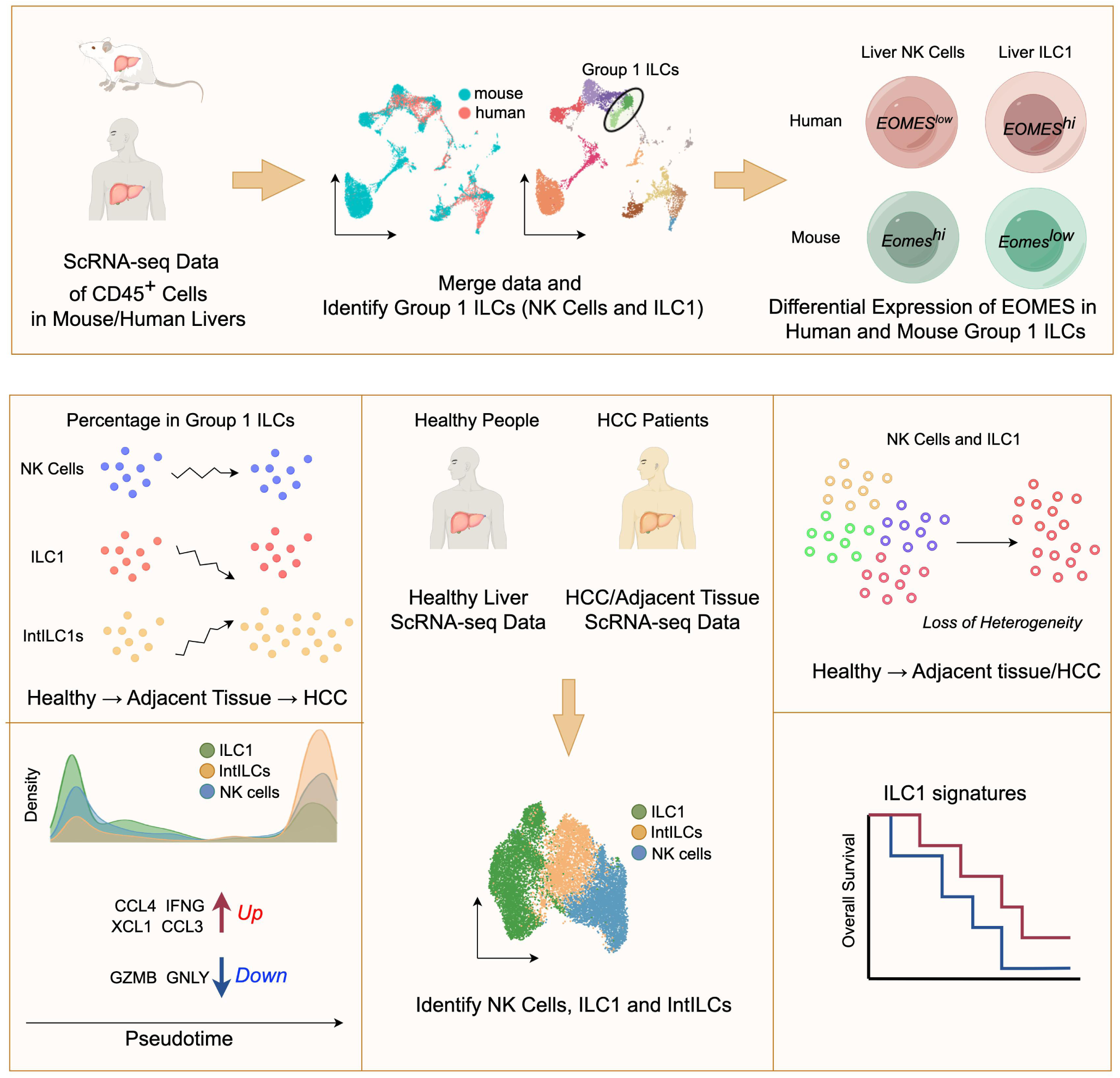

## Introduction

The liver is one of the most important digestive organs, generating innate and adaptive immune responses against pathogen, malignant transformed cells, and harmful molecules[1–3]. Thanks to supplied by arterial and venous blood, liver is also an organ prone to tumorigenesis and a common target for malignant metastasis[4]. Based on the type of cells that the cancer initiated from, there are two different type of liver cancer: hepatocellular carcinoma (HCC) and intrahepatic cholangiocarcinoma[5, 6]. HCC is the most common primary liver cancer[7]. A variety of immune cells, which including macrophages, natural killer (NK) cells, NKT cells, dendritic cells (DCs), eosinophils, δγ T cells and conventional T cells were discovered in the liver[8–10]. Liver immune cells eliminate pathogens, maintain liver immune homeostasis, and monitor malignant tumor cells[11, 12]. Group(Type) I ILCs is an innate population of liver immune cells, consisting of conventional NK cells and ILC1s[13]. Group I ILCs account for about 30% of the total number of human intrahepatic lymphocytes[14], 15 ∼ 40% for mouse liver lymphocytes[15, 16]. NK cells recognize and eliminate virus infected or malignant transformed tumor cells[17–20]. ILC1s has been shown to be a distinct lineage from NK cells in mouse livers[21]. One of the important pathways through which ILC1s work is the secretion of cytokines, mainly IFN-γ. ILC1s can protect the host from mouse cytomegalovirus (MCMV) and bacterial at the sites of infection[22–24]. ILC1s limit viral replication and promote host survival prior to the recruitment of lymphocytes into infected tissue by rapidly producing IFN-γ after DCs activation and interleukin (IL)-12 production[22]. When ILC1s accumulate in inflamed tissue, they may exacerbate fibrosis and tumor growth by secreting TGF- [25].

Mouse liver group I ILCs, which include NK cells and ILC1, are characterized as CD45^+^CD3^-^NK1.1^+^NKp46^+^. These cells represent distinct development lineages. The differential expression of CD49a and CD49b is employed to distinguish NK cells (CD49a^-^CD49b^+^) from ILC1 (CD49a^-^CD49b^+^) via flow cytometry at homeostatic state[15, 21]. In addition, functional differences were also observed in NK cells and ILC1s. ILC1s have lower in vitro killing effect on tumor cells and expressed less cytotoxic molecules granzyme A/B and perforin compared with NK cell in liver[26]. However, ILC1 in mammary tumor model of mice expressed more granzyme C and TRAIL than NK cell, both of which could induce apoptosis of target cells[27]. Although the studies on mouse liver ILC1s have made rapid progress, there are still plenty of unsolved questions in the identification, development, function, and regulation of human liver ILC1s. Studies have shown that similar to mouse liver ILC1, human liver NK cells (CD3^-^CD56^+^NKp46^+^) can also be divided into two subpopulations based on the expression of CD49a; In comparison to conventional NK cells, CD49a^+^ human intrahepatic NK cells, which are found within the liver, have been found to produce higher level of proinflammatory cytokines, and are absent of the transcriptional factor Eomes[28]. Although both mouse NK cells and ILC1 are T-bet dependent[26, 29], some other studies showed that there is a CXCR6^+^ tissue resident population of human liver NK cells, which are Eomes^+^T-bet^-^[30, 31]. It is uncertain whether CXCR6^+^Eomes^+^T-bet^-^ NK cells in human liver fully resemble hepatic ILC1s in mouse.

Signel-cell RNA sequencing (scRNA-seq) is a powerful technique that allows for the detailed analysis of the transcriptome (gene expression) at the single cell level[32]. It has been extensively used to study immune cell populations, track changes in gene expression during development and disease progression[33–35]. In this study, we utilized scRNA-seq data from public databases to conduct a detailed analysis of human liver group I ILCs, with a particular focus on healthy individuals and patients with HCC. Our preliminary data-driven analysis successfully identified a subset of cells in the human liver that closely resemble mouse liver ILC1s. Our study highlights the variable expression of *EOMES* across human and mouse models, suggesting species-specific regulatory mechanisms. Additionally, we discovered and characterized a distinct cluster of CD127^+^ intermediate ILCs that display characteristics of both NK cells and ILC1s. This finding provides a unique perspective on immune regulation within the liver. Through comprehensive characterization of these cell groups and investigation into their dynamic alterations during tumor progression, our research enhances understanding of the hepatic tumor immune environment. Insights gained from this study could potentially inform new strategies for the prevention and treatment of liver cancer, emphasizing the clinical relevance of our findings.

## Materials and Methods

### ScRNA-seq data processing and analysis

Our study used a total of 15,824 liver CD45^+^ cells from mice and humans (6,038 from humans and 9,786 from mice) and 12,526 group I ILCs from healthy human and HCC livers (6,008 cells from healthy liver, 3939 cells from HCC and 2,579 cells from adjacent normal tissue). Gene expression matrix data were downloaded from GEO(Gene Expression Omnibus) public database at NCBI. Firstly, the data set of each sample was subject to quality control, and cells were screened according to the two indexes of gene number and mitochondrial UMI. The Seurat package was then used to normalize the gene matrix for each batch separately. The normalized gene matrix was converted to Z-scores using the ScaleData function implemented in the Seurat package, and the data from the different batches were integrated using the canonical correlation analysis (CCA) method in Seurat. We used the top 2,000 highly variable genes as inputs to run PCA, and then selected the first 20 principal components for UMAP visualization and clustering (the first 15 PC for human healthy and HCC liver data). Downstream data processing and analysis steps, including filtering, normalization, PCA, clustering, UMAP, and differential gene expression analyses, were performed using the Seurat package version 4.0.1. We also used clusterProfiler R package v4.11.0 for enrichment analysis (GSEA, KEGG and GO analysis) and Monocle R package v2.30.0 for pseudotime analysis.

We annotated cell clusters based on immune cell-specific marker genes: Group I ILCs (mouse:*Ncr1*:human:*KLRF1*), B cells(*CD79A, MS4A1*), Plasma cells(*JCHAIN, MZB1*), CD8^+^ T cells(*CD3D, CD8A*), CD4^+^ T cells(*CD3D, CD4, IL7R*), Monocytes(*LYZ, VCAN*), Macrophages(*CST3, CD68*), Kupffer cells(*C1QA, C1QB, CLEC4F*), Neutrophils(*CSF3R, PTGS2*), NKT cells(*CD3E, GZMB*), Epithelial cells(*AQP1, SEPP1*) (Figure S1).

The Cancer Genome Atlas (TCGA) data were obtained, analyzed, and plotted using the public database: Gene Expression Profiling Interactive Analysis database (GEPIA2, http://gepia2.cancer-pku.cn/#index) [36]. The ILC1 signature was defined as the mean normalized expression (log2 (TPM + 1)) of marker genes included *XCL1, CD160, CCL3, EOMES, IL2RB, CXCR6, XCL2, KLRB1, KLRC1*. Samples were divided into high and low expression group by median value. Difference in overall survival time was showed in Kaplan–Meier survival curves, and statistical significance was determined by log-rank p values.

## Results

### Mapping the Similarities between Human and Mouse Group **I** ILCs

To construct a comprehensive atlas of human NK cells spanning both healthy and cancerous liver tissues, we utilized six published scRNA-seq datasets encompassing NK cells derived from both healthy and cancer-affected liver tissues (Figure 1A). We additionally obtained two mouse scRNA-seq datasets from healthy liver tissues to serve as controls, facilitating a comparative analysis of the relationship between mouse and human liver group I ILCs (Figure 1A). Datasets of human and mouse healthy liver tissues were initially integrated, setting the foundation for a more precise identification of human liver conventional NK (cNK) cells and ILC1s.

**Figure. 1.**
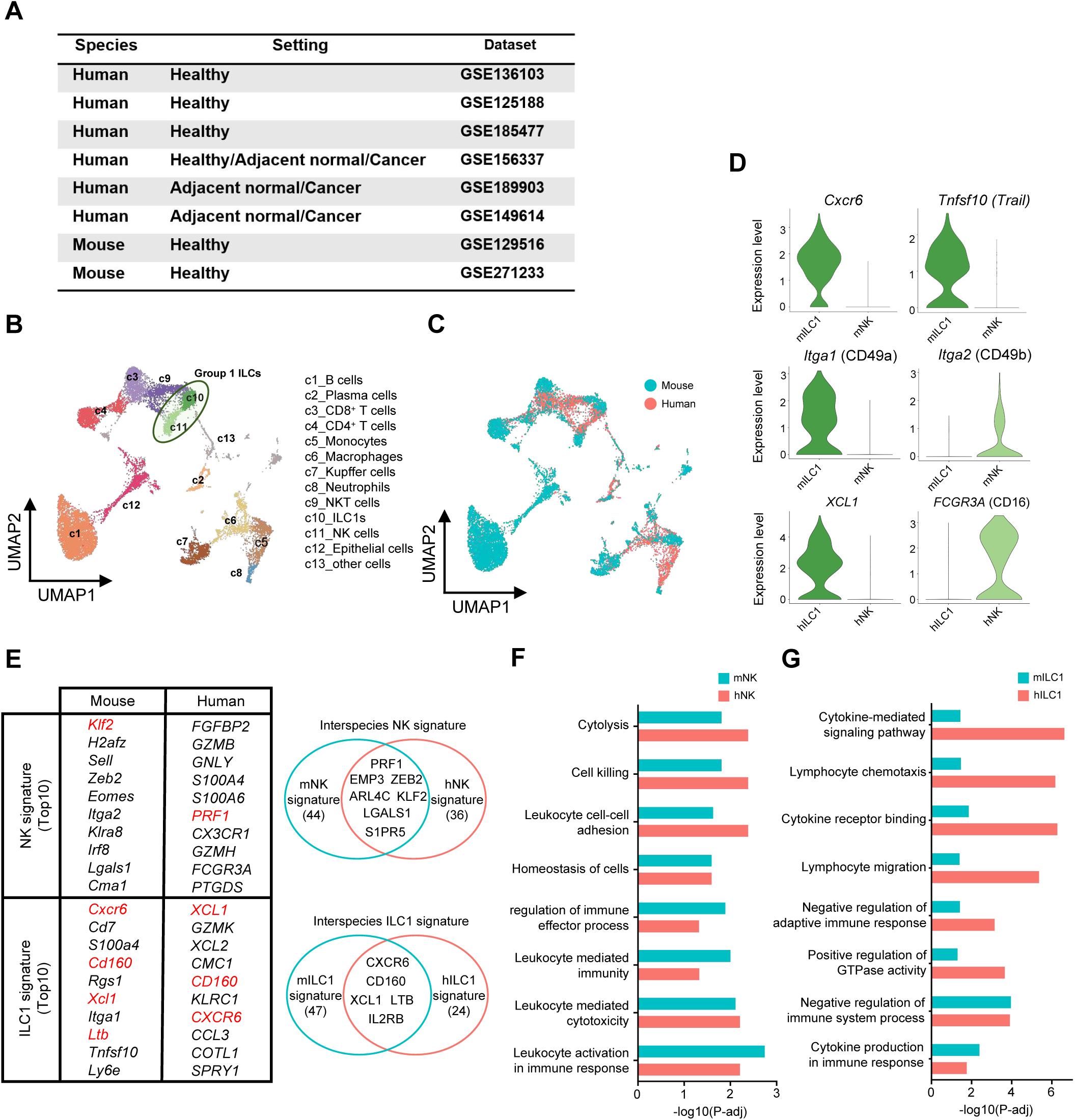
A unique human liver ILC1 population identified with reference to mouse liver ILC1. **(A)** Table of all publicly available data set information used in our study. **(B)** UMAP plot of all the 15,824 CD45^+^ cells from 2 mouse liver datasets and 4 human liver datasets. Cells are colored based on the 13 clusters defined by clustering and marker genes. **(C)** UMAP plot of all the CD45^+^ cells colored by sample sources, human (red), mouse (blue). **(D)** Violin plots of expression levels of selected markers between NK cells and ILC1 (*Cxcr6, Tnfsf10, Itga1* and *Itga2* expression in mouse liver, *XCL1* and *FCGR3A* expression in human liver). **(E)** Table and venn diagram of differential genes in human and mouse liver NK cells and ILC1. Venn diagram shows the interspecies signature. **(F and G)** Enriched Gene Ontology (GO) terms of DEGs in NK cells and ILC1s. The Enrichment gene set in upregulated genes of mNK (blue) and hNK (red) **(F)** mILC1 (blue) and hILC1 (red) **(G)** were indicated in different colours. Bar length represents statistical significance.

Through random sampling and normalization, we obtained 15,824 cells (6,038 from humans and 9,786 from mice) for analysis (Figure 1B). UMAP (Uniform Manifold Approximation and Projection) analysis of scRNA-seq data from mouse and human liver cells identified 12 clearly defined cell clusters based on the expression of canonical marker genes, including *C1QA*, *C1QB*, and *CLEC4F* for Kupffer cells, *CD79A* for B cells, *CD3D* and *CD8B* for CD8^+^ T cells, and *GNLY* for NK cells (Table S1; Figure S1)[37]. Of note, most of clusters were largely overlapping between human and mouse (Figure 1C).

Flow cytometry was used to differentiate mouse NK cells and ILC1s. The phenotype of mouse NK cells was found to be CD49a^-^CD49b^+^TRAIL^-^CXCR6^-^Eome^+^NKp46*^+^* phenotype, while that of ILC1s was determined to be CD49a^+^ CD49b^-^ TRAIL^+^ CXCR6^+^ Eomes^-^ NKp46^+^ phenotype[38]. The differentiation of NK cells from ILC1s in mice can be achieved by using marker genes such as *Itga1, Itga2, Trail, Cxcr6,* and *Eomes*[39, 40]. In this study, we applied scRNA-seq data to delineate mouse liver NK cells and ILC1s, utilizing specific markers - namely *Itga1*, *Itga2*, *Tnfsf10* (*Trail*), and *Cxcr6* to achieve this differentiation (Figure 1D). Through integrated analysis and cross-clustering of human and mouse samples, human liver NK cells and cells comparable to mouse liver ILC1s were identified. Notably, counterparts to mouse liver NK cells and ILC1s were consistently detected in all three healthy human liver *CD45^+^* cell samples, indicating that ILC1s were a stable group of cells in the human liver. For ease of reference, mouse liver NK cells and ILC1s were termed mNK and mILC1s, respectively, while their human equivalents were designated as hNK and hILC1s.

To further confirm the similarity between hILC1s and mILC1s, signatures of NK cells and ILC1s were compared between humans and mice. Both mNK and hNK cells exhibit high expression of *PRF1*, *EMP3*, *ZEB2*, *ARL4C*, *KLF2*, *LGALS1*, and *S1PR5*, while *CXCR6*, *CD160*, *XCL1*, *LTB*, and *IL2RB* serve as key markers across mILC1 and hILC1 (Figure 1E). Gene Ontology (GO) assessments were conducted to elucidate the functional characteristics of NK cells and ILC1s. The outcomes demonstrated that genes highly expressed in both hNK and mNK cells were predominantly involved in cytolysis, leukocyte-mediated cytotoxicity, and cell killing, indicating a marked similarity in their capacities for these effector functions compared to hILC1s or mILC1s, respectively (Figure 1F). Compared with NK cells, hILC1s and mILC1s mainly involved in migration, chemotaxis, and cytokine-mediated signaling pathways, indicating the role of ILC1s in immune regulation (Figure 1G). Notably, hILC1s exhibit more pronounced activity in chemokine and cytokine pathways than mILC1 cells, demonstrating the functional diversity between these cell types.

### Differential Expression of EOMES in Human and Mouse Group **I** ILCs

Eomes plays a crucial role in the differentiation and homeostatic maintenance of mNK and mILC1 cells in the mouse liver. The development of mNK cells depends on Eomes, and high expression level of Eomes is also considered as one of the hallmarks of mNK cells[29]. Previous studies have found that EOMES^hi^ T-bet^lo^ CD56^bright^ NK cells are enriched in the human liver, representing a subset of liver-resident NK cells with increased tolerance [31]. Although there has been significant research into the role of EOMES in human NK (hNK) cells, these studies have predominantly relied on NK cells isolated from peripheral blood or derived via in vitro differentiation from stem cells. As a result, they may not accurately reflect the function of EOMES in liver-resident NK cells or hepatic ILC1s [41]. In mice, the expression level of *Eomes* in liver mNK cells is much higher than that in mILC1s (Figure 2A). In contrast, in human liver, the expression of *Eomes* in hNK cells is significantly lower than in hILC1s (Figure 2A). This discrepancy not only highlights the differences in transcription factor regulation between human and mouse group I ILCs but also raises questions about using *EOMES* to distinguish or define human liver hNK and hILC1 cells.

**Figure. 2.**
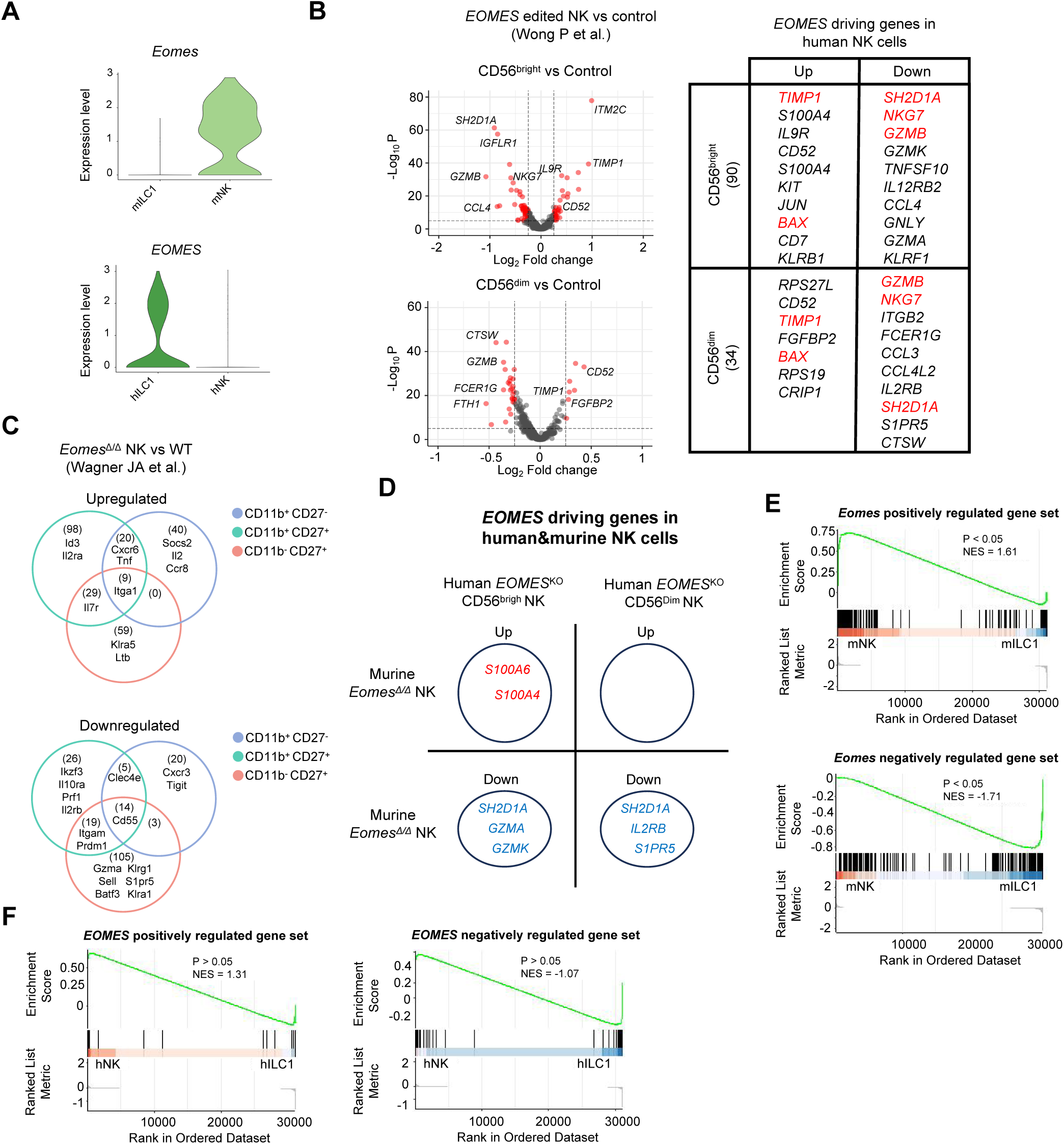
EOMES in hILC1 and mNK have different transcription factor regulatory networks. **(A)** Violin plots of EOMES expression in mouse and human liver NK cells and ILC1s. **(B)** Volcano plot and table of differential genes in EOMES knockout human liver CD56+ NK cells and control (GSE227636). There are two groups of CD56bright NK cells and CD56dim NK cells. **(C)** Venn diagram of EOMES knockout mouse spleen NK cells and control (GSE132942). There are three groups of CD11b+ CD27-, CD11b+ CD27- and CD11b+ CD27- NK cells. **(D)** Overlapped EOMES driving genes in human and murine NK cells. **(E)** Gene Set Enrichment Analysis (GSEA) showing the enrichment of *Eomes* positively regulated gene set and *Eomes* negatively regulated gene set of DEGs in mNK compared mILC1. **(F)** Gene Set Enrichment Analysis (GSEA) showing the enrichment of *EOMES* positively regulated gene set and *EOMES* negatively regulated gene set of DEGs in hNK compared hILC1. NES, normalized enrichment score. p, P-value.

A pair of separate studies employed different genetic editing techniques to knockout the *EOMES* gene in NK cells. CRISPR was used to knockout *EOMES* in human NK cells, resulting in an increase in the expression of *TIMP1* and *BAX*, and a decrease in the expression of *SH2D1A*, *NKG7*, and *GZMB* (granzyme B) (Figure 2C) [42]. Meanwhile, a conditional knockout technique was applied in mouse NK cells, leading to the observation that 9 genes showed increased expression at all three stages of NK cell development (e.g. *Itga1*), while 14 genes showed decreased expression (e.g. *Cd55*). Only *S100A6* and *S100A8* were upregulated in both human and mouse NK cells, while *SH2D1A*, *GZMA*, *GZMK*, *IL2RB*, and *S1PR5* were commonly downregulated (Figures 2C and D)[43].

To further investigate the role of *Eomes* in NK and ILC1 cells, we utilized data from *Eomes* conditional knockout mice (GSE132942) to identify target genes of *Eomes*, thereby studying its functions in NK and ILC1 cells[43]. Following the knockout of *Eomes*, genes that were downregulated in NK cells were defined as the *Eomes* positively regulated gene set, while genes that were upregulated were defined as the *Eomes* negatively regulated gene set. These two gene sets were subsequently subjected to GSEA in mNK and mILC1 cells. Consistent with the elevated expression of *Eomes* in mNK cells, the *Eomes* positively regulated gene set was significantly enriched in mNK cells, whereas the *Eomes* negatively regulated gene set was significantly enriched in mILC1 cells (Figures 2F). These findings indicate that *Eomes* positively regulated genes are more active in mNK cells, while *Eomes* negatively regulated genes are more active in mILC1 cells, highlighting the distinct regulatory roles of *Eomes* in these cell types.

To extend these findings to human data, we employed data derived from *EOMES* knockout human CD56^+^ cells (Pamela and Todd, GSE227636)[42] to identify specific target genes of *EOMES* within hNK cells and hILC1. Distinct from murine models, where genes upregulated by *Eomes* are predominantly enriched in mNK cells expressing high levels of *Eomes*, and genes downregulated by *Eomes* are enriched in mILC1 cells with low *Eomes* expression, the human data presents a contrasting scenario. In hILC1 cells, despite high expression of *EOMES*, the genes that were highly expressed, when compared to hNK cells, did not show significant enrichment in the EOMES positively regulated gene set. Similarly, in hNK cells with low *EOMES* expression, the genes downregulated by *EOMES* were not significantly enriched either (Figures 2F). This discrepancy highlights fundamental differences in *EOMES* expression patterns between murine and human hepatic group I innate lymphoid cells, suggesting that the regulatory role of *EOMES* in human liver hNK and hILC1 cells may not be as critical as observed in murine models. These findings also underscore the necessity of specific approaches when studying the functionality of human liver group I ILCs, indicating clear divergences in transcriptional regulation and cellular function between species.

### Gene Expression Profiles of Group **I** ILCs in HCC

To further investigate group L innate lymphoid cells in the liver immune microenvironment of hepatocellular carcinoma (HCC), we analyzed NK cells and ILC1s from 6 published human scRNA-seq datasets, which encompass both liver and HCC tissues (Figure 3A). A total of 12,526 group L ILCs (6008 cells in healthy liver, 2,579 cells in adjacent tissue, and 3,939 cells in HCC) were included in this analysis. Using the UMAP method, three major clusters of group I ILCs were identified. Beyond the established NK and ILC1 clusters, a distinct cluster characterized by moderate expression of NK/ILC1 markers was observed, which was defined as intermediate ILCs (intILCs). Notably, clustering patterns specific to tumor and adjacent tissues were revealed by this analysis, suggesting potential cellular remodeling within the HCC microenvironment (Figure 3A and B).

**Figure. 3.**
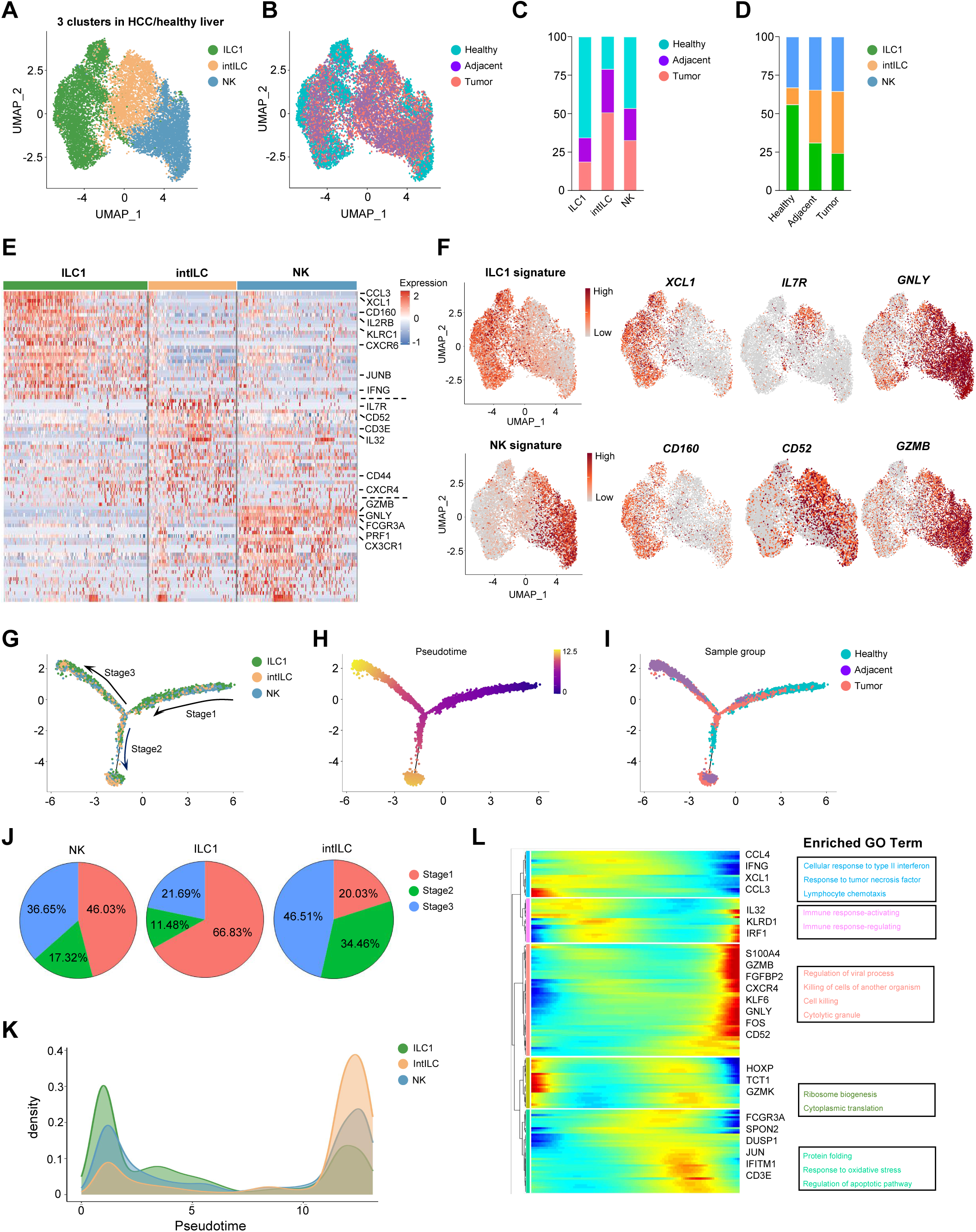
Global changes in NK cells and ILC1 in hepatocellular carcinoma. (A and. **B)** UMAP plot of type 1 ILCs in HCC, adjacent normal tissue and healthy liver colored by 3 clusters, ILC1 (green), intILC (yellow) and NK cells (blue) **(A)** and colored by 3 sample groups, Healthy (lightblue), Adjacent (purple) and Tumor (red) **(B)**. **(C)** Percentages of cells in Healthy, Adjacent and Tumor samples in 3 clusters. **(D)** Percentages of NK cells, intILC and ILC1s in 3 sample groups. **(E)** Heatmap of the top-ranking cluster specific genes. Columns denote cells; rows denote genes. **(F)** UMAP plot showing the expression levels of NK/ILC1 signature and some cell marker of 3 clusters, *XCL1, IL7R, GNLY, CD160, CD52* and *GZMB*, across type 1 ILCs. **(G-I)** The ordering of different cell clusters along pseudotime in a two-dimensional state-space defined by Monocle2. Cell orders were inferred from the expression of the differential genes across sample groups. Each point corresponds to a single cell, and cells are colored by clusters **(G)**, pseudotime value **(H)** and sample groups **(I)**. **(J)** Sector graph of the proportions of NK cells, ILC1s, and intILC in the three stages of the pseudo-timing. **(K)** Plot of cell density along pseudotime of ILC1s, intILC and NK cells. **(L)** Heatmap showing cell-state transitions related genes which were divided into five clusters in Monocle2 analysis (left) and enriched Gene Ontology (GO) terms of the genes of five clusters (right). The depth of the color from blue to red represents the expression level from low to high.

In the healthy liver, ILC1 cells represent the predominant population, accounting for more than 50% of group L ILCs. NK cells are the next most abundant, making up approximately 30% of group L ILCs in the liver. intILCs constitute about 15% of the population. Across the progression from healthy liver to tumor-adjacent tissue and onto tumor tissue, the proportion of NK cells remains relatively stable. In contrast, the proportion of ILC1 cells gradually decreases, whereas that of intILCs progressively increases (Figure 3, C and D). The heatmap illustrated variations in gene expression among the clusters: ILC1s predominantly expressed *CCL3*, *XCL1*, and *CD160*; NK cells showed elevated levels of *GZMB*, *GNLY*, and *FCGR3A*; and intILCs were characterized by higher levels of *IL7R*, *CD52*, *CD3E*, *CD44*, and *CXCR4* (Figure 3E). Intermediate levels of both NK and ILC1 marker genes were detected in intILCs.

Group L ILCs trajectories and dynamics were analyzed using pseudotime to predict the characteristic of cells in distinct states (Figure 3, G-I). A clear directional flow was shown in the trajectory, containing two termini corresponding to different cell fates. The root of the trajectory was mainly populated by cells from healthy samples, while the two termini were mainly primarily by cells from tumor and adjacent tissues (Figure 3, G and H; Figure S2, A). As cancer progressed, the proportion of intILCs increased significantly, with a notable decrease in the ILC1 proportion, from adjacent tissues to tumor (Figure 3, I-K). Next, we analyzed the DEGs in different stages of cell fates to explore the potential signals and gene patterns during tumor progression (Figure 3L; Figure S2, D).

Cytokine genes including *CCL4*, *IFNG*, *XCL1*, and *CCL3* showed a progressive down-regulation from healthy to tumor tissues (Figure 3L). Simultaneously, functional genes such as *GZMB* and *GNLY* in NK cells exhibited an incremental up-regulation (Figure S2, B,C). In contrast, the expression profiles of intILC and NK cell markers increased in correlation with cell fate progression, suggesting a potential shift from ILC1 to intILC during cancer advancement (Figure 3L). We observed that the trajectory branch containing healthy components was positively associated with cytokine production and immune response, while the branches with adjacent normal and tumor components exhibited elevated cytotoxicity scores (Figure 3L). These trajectory data suggest significant plasticity and aberrant cellular states in the ILC1-intILC-NK subsets during HCC progression.

### Subpopulation Dynamics of NK Cells and ILC1s in HCC

250 significant differentially expressed genes (DEGs) with log2 |fold change| > 0.5 and P adj < 0.05 were identified in ILC1s from tumor and adjacent liver tissues compared to healthy controls (Figure 4G). Compared to healthy liver ILC1s, global transcription changes in ILC1s from tumor and adjacent tissues were highly correlated, indicating similar regulatory alterations (Figure 4G). Relative to healthy samples, 118 DEGs, including key ILC1 markers such as *CCL3*, *XCL1*, *CD160*, *EOMES*, *IL2RB*, *NFKBIA*, and *JUNB*, were downregulated in both tumor and adjacent tissues (Figure 4G). This reduction in marker gene expression levels suggests that a transformation of liver ILC1s into other cell types may be occurring during HCC progression. Furthermore, increased expression of 131 genes, including *HSPA1A*, *CREM*, *CXCR4*, *TNFAIP3*, *ZEB2*, and *STAT4*, was observed in both HCC and adjacent tissues (Figure 4G). In ILC1s from these tissues, genes upregulated were predominantly involved in pathways related to oxidative stress and apoptosis, while those associated with leukocyte-mediated cytotoxicity and cell killing were found to be downregulated in both tumor and adjacent liver samples (Figures 4H and I).

**Figure. 4.**
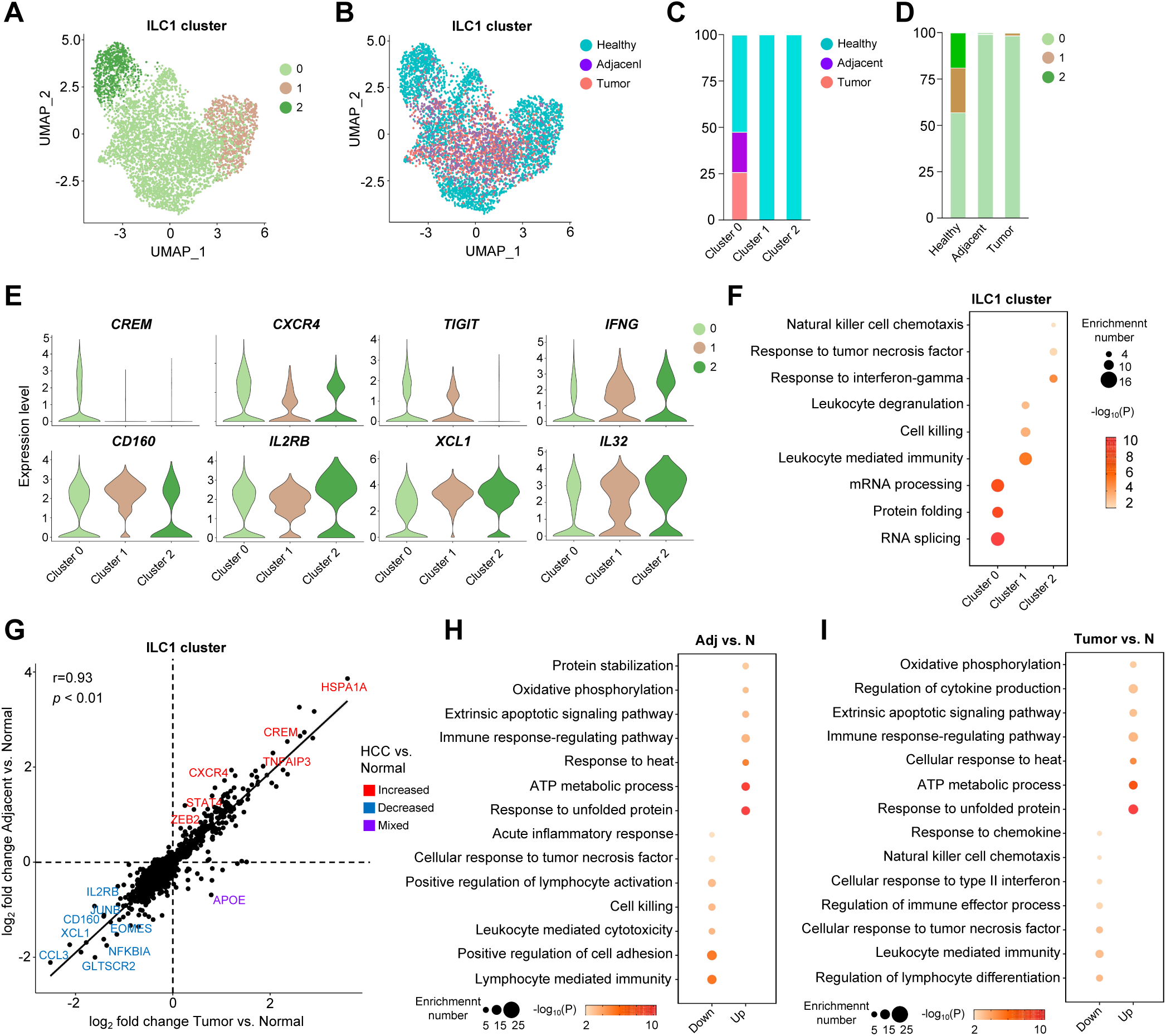
scRNA-seq reveals unique properties of ILC1s in the tumor environment. (A and. **B)** UMAP plot of ILC1 subsets in HCC, adjacent normal tissue and healthy liver. Cells colored by 3 subclusters **(A)** and by 3 sample groups**(B)**. **(C)** Percentages of cells in Healthy, Adjacent and Tumor samples in 3 subclusters. **(D)** Percentages of 3 subclusters in HCC, adjacent normal tissue and healthy liver. **(E)** Violin plots showing the expression of some selected genes in 3 subclusters, *CREM, CXCR4, TIGIT, IFNG, CD160, IL2RB, XCL1* and *IL32*. **(F)** Enriched GO term of marker genes in 3 subclusters. Dot size represents enriched gene number, and color intensity represents significance. **(G)** Pearson’s correlation of the log_2_ fold change of DEGs between tumor and normal samples, and adjacent normal and normal samples. Labels show the selected significant increased (red), decreased (blue), and mixed (purple; increased in tumor samples and decreased in adjacent samples or vice versa) DEGs in tumor or adjacent normal samples. **(H and I)** Enriched GO term of DEGs in ILC1s from Adjacent compared ILC1s from Normal **(H)** and in ILC1s from Tumor **(I)** compared ILC1s from Normal. Dot size represents enriched gene number, and color intensity represents significance.

Subpopulation analysis using scRNA-seq offers an unprecedented resolution to characterize the heterogeneity of cell populations, enabling a more thorough examination of differences in gene expression, metabolic pathways, and cell function within distinct cellular subgroups. In this study, we separately selected NK cells, ILC1s, and the newly identified intILCs for detailed subpopulation analysis to explore their characteristics and examine their changes within HCC. ILC1s were divided into three clusters (Figure 4A), with distinct distributions among each sample (Figure 4B). Three distinct ILC1 subgroups are well-represented in normal liver tissue, comprising 56.9% in cluster 0, 24.1% in cluster 1, and 19% in cluster 2 (Figure 4, C and D). In contrast, ILC1s in HCC samples and adjacent cancer tissues are predominantly confined to cluster 0, indicating a significant shift in subgroup composition in the disease state.

Cluster 0 ILC1 displayed high expression levels of *CREM*, *CXCR4*, and *TIGIT*, similar to the exhausted NK cells that were previously reported in cancer patients[44] (Figure 4E). Clusters 2 and 3 ILC1 were characterized by *IFNG*, *CD160*, *IL2RB*, *XCL1*, and *IL32*, showing similar expression of ILC1 markers (Figure 4E). GO analysis was used to further identify the potential functions of different ILC1 clusters. Cluster 0 showed high translation activity compared to other clusters (Figure 4F). Cell-killing related genes were enriched in cluster 1, indicating the potential cytotoxicity of this cluster (Figure 4F). The ability to response to cytokine like IFN-γ and TNF-α was active in cluster 2 ILC1 (Figure 4F). These data suggest that clusters 1 and 2 ILC1 consist of populations with cytotoxicity or cytokine response function, while cluster 0 might contain exhausted ILC1s with suppressed anti-tumor immunity. Tumors may evade immune surveillance by inducing normal ILC1s to adopt exhausted phenotypes.

*CREM*, *TNFAIP3*, *CXCR4*, *STAT4*, *CCL5*, and *ATF3* was upregulated in NK cells from HCC patients, while *CCL3*, *MAF1*, *PTPN6*, *XCL1*, *CD48*, and *ID2* was downregulated (Figure 5G). GO analysis revealed enrichment of ATP metabolic process, cellular response to heat, response to unfolded protein, and oxidative phosphorylation in upregulated DEGs (Figure 5, H and I). Pathways associated with lymphocyte mediated immunity, regulation of immune effector process, natural killer cell chemotaxis, and wound healing were enriched in downregulated DEGs (Figure 5, H and I). Sub-clustering of NK cells revealed three distinct clusters (Figure 5A). In healthy liver samples, cluster 0 contained 62% of NK cells, with the remainder in cluster 1 (21.9%) and cluster 2 (15.8%) (Figure 5D). In HCC tumors and adjacent liver tissues, cluster 0 was predominant, encompassing almost all NK cells, with clusters 2 and 3 being virtually absent (Figure 5, B and C). NK cells in cluster 0 were characterized by high levels of *PTGDS*, activation marker CD69, and cytotoxic protein *GZMB*, signifying strong tumor-killing capabilities (Figure 5E). NK cells in cluster 1 expressed *XCL1* and *CD160*, showing traits akin to ILC1s (Figure 5E). Cluster 2 NK cells exhibited high levels of C-C motif chemokine ligand 4 (*CCL4*) and JUNB, with only subtle differences from cluster 1 (Figure 5E). Both clusters 1 and 2 demonstrated responsiveness to interferon, but cluster 2 additionally showed enhanced migration abilities and activated ERK signaling (Figure 5F).

**Figure. 5.**
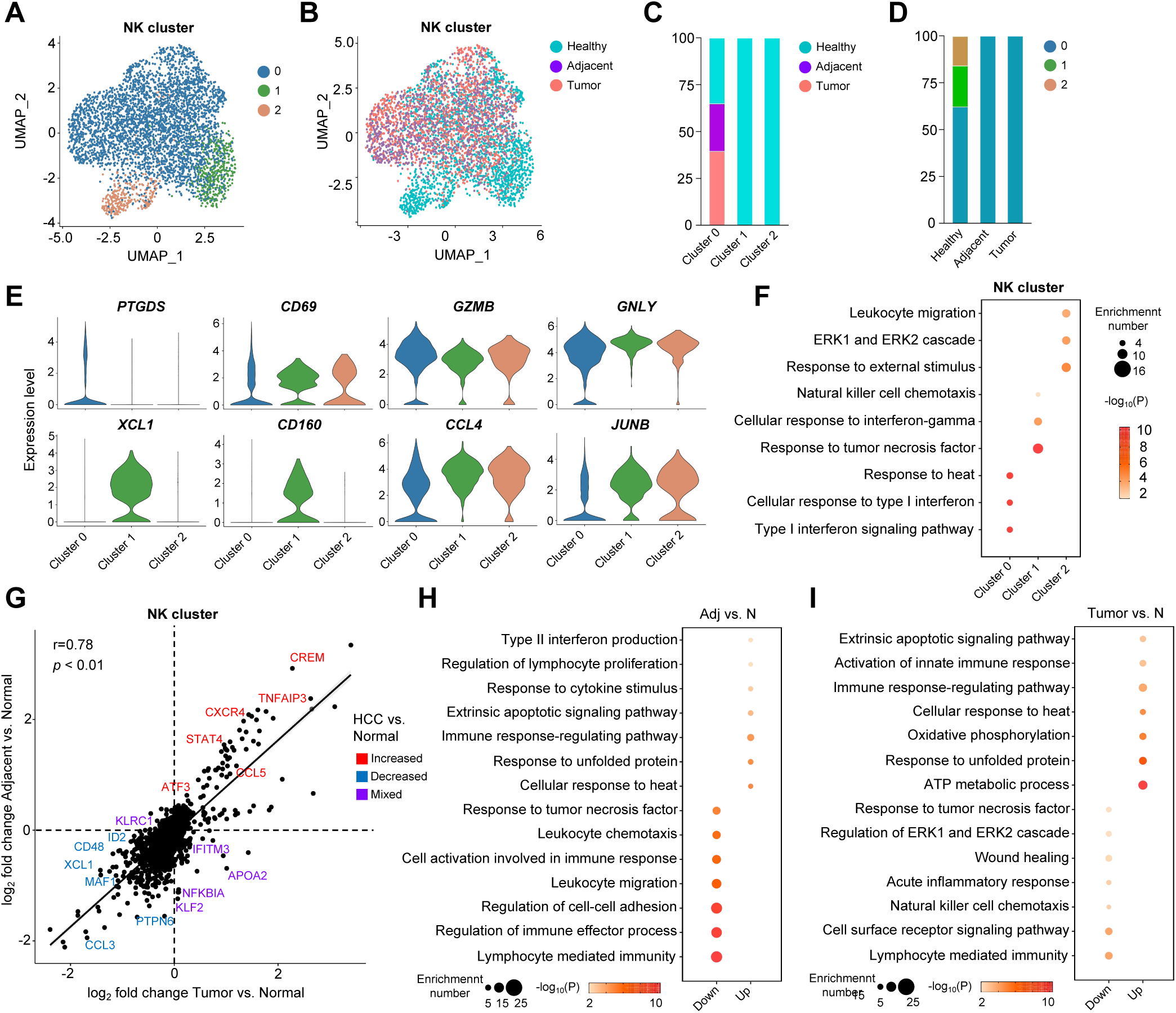
scRNA-seq reveals unique properties of NK cells in the tumor environment (A and. **B)** UMAP plot of NK cells subsets in HCC, adjacent normal tissue and healthy liver. Cells colored by 3 subclusters **(A)** and by 3 sample groups **(B)**. **(C)** Percentages of cells in Healthy, Adjacent and Tumor samples in 3 subclusters. **(D)** Percentages of 3 subclusters in HCC, adjacent normal tissue and healthy liver. **(E)** Violin plots showing the expression of some selected genes (*PTGDS, CD69, GZMB, GNLY, XCL1, CD160, CCL4* and *JUNB*) in 3 subclusters. **(F)** Enriched GO term of marker genes in 3 subclusters. Dot size represents enriched gene number, and color intensity represents significance. **(G)** Pearson’s correlation of the log_2_ fold change of DEGs between tumor and normal samples, and adjacent normal and normal samples. Labels show the selected significant increased (red), decreased (blue), and mixed (purple; increased in tumor samples and decreased in adjacent samples or vice versa) DEGs in tumor or adjacent normal samples. **(H and I)** Enriched GO term of DEGs in NK cells from Adjacent compared NK cells from Normal **(H)** and in NK cells from Tumor compared NK cells from Normal **(I)**. Dot size represents enriched gene number, and color intensity represents significance.

### Identification of different Tumor-Associated intermediate ILC Populations

One of the pivotal findings of this study was the identification of intILCs and their elevated prevalence in both tumor and adjacent liver tissues. Similar to NK cells and ILC1s, the genes *CCL3*, *PTGDS*, *FCER1G*, *KLRB1*, and *CD160* were found to be downregulated in both tumor tissues and adjacent liver regions. In contrast, *CREM*, *TNFAIP3*, *ANXA1*, and *CXCR4* showed upregulation (Figure 6I). The GO analysis of differentially expressed genes in intILCs revealed consistent biological functions: lymphocyte-mediated immunity and the cell surface receptor signaling pathway were significantly downregulated in tumor and adjacent tissues. Meanwhile, processes such as ATP metabolism, cellular response to heat, and oxidative phosphorylation were enhanced (Figure 6, J and K). These findings indicate that the tumor microenvironment may also trigger specific stress responses and metabolic shifts, or metabolic reprogramming, in intILCs.

**Figure. 6.**
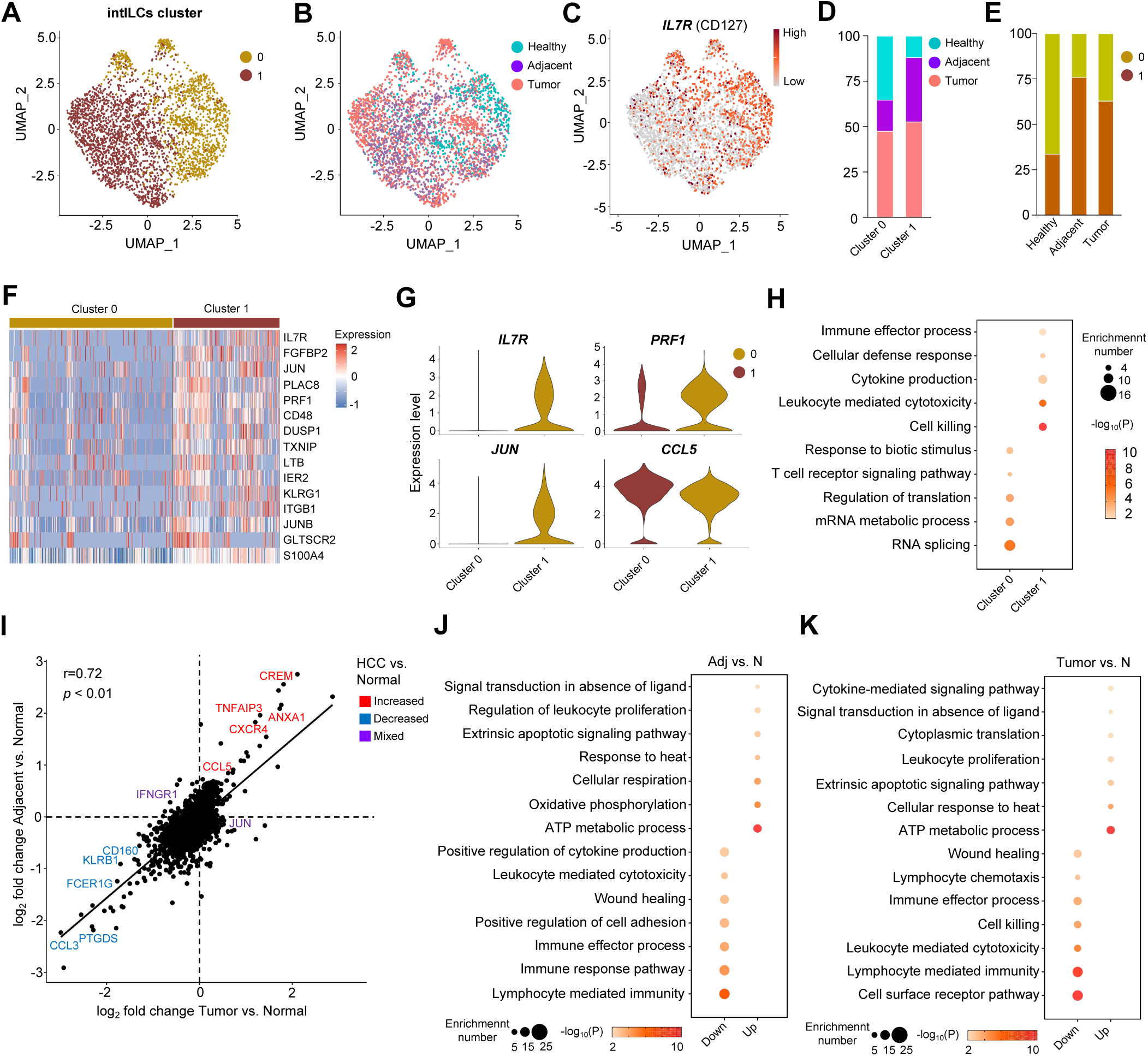
scRNA-seq reveals unique properties of Tumor-Associated intermediate ILC Populations. (A and. **B)** UMAP plot of intILCs subsets in HCC, adjacent normal tissue and healthy liver. Cells colored by 2 subclusters **(A)** and by 3 sample groups**(B)**. **(C)** UMAP plot showing the expression of *IL7R (CD127)* in intILCs. **(D)** Percentages of cells in Healthy, Adjacent and Tumor samples in 2 subclusters. **(E)** Percentages of 2 subclusters in HCC, adjacent normal tissue and healthy liver. **(F)** Heatmap of the 15 specific genes in cluster 1. Columns denote cells; rows denote genes. **(G)** Violin plots showing the expression of some selected genes in subclusters, *IL7R, PRF1, JUN* and *CCL5*. **(H)** Enriched GO term of marker genes in 2 subclusters. Dot size represents enriched gene number, and color intensity represents significance. **(I)** Pearson’s correlation of the log_2_ fold change of DEGs between tumor and normal samples, and adjacent normal and normal samples. Labels show the selected significant increased (red), decreased (blue), and mixed (purple; increased in tumor samples and decreased in adjacent samples or vice versa) DEGs in tumor or adjacent normal samples. **(J and K)** Enriched GO term of DEGs in intILCs from Adjacent compared intILCs from Normal **(J)** and in intILCs from Tumor compared intILCs from Normal **(K)**. Dot size represents enriched gene number, and color intensity represents significance.

UMAP analysis further segmented these intILCs into two distinct clusters (Figure 6, A and B), which were prominently present in healthy liver, tumor, and tumor-adjacent tissues. A primary distinguishing marker between these clusters is *CD127*, the IL-7 receptor (Figure 6C). In the healthy liver, cluster 0 (*IL7R*^hi^) accounted for approximately 70% of intILCs. Conversely, in both tumor tissues and adjacent liver tissues, there was a noticeable rise in cluster 1 (*IL7R*^lo^), where it comprised over 50% of the intILCs (Figure 6, D and E). *IL7R*^hi^ intILCs demonstrated a distinctive gene expression profile compared to IL7R^lo^ intILCs, including upregulation of *FGFBP2*, *JUN*, *PRF1*, *CD48*, *CCL5*, among others (Figure 6, F and G). Compared to their *IL7R*^lo^ counterparts, *IL7R*^hi^ intILCs exhibited enhanced signaling related to cytotoxicity, cytokine production, and cellular defense responses (Figure 6H). This pattern suggests a reduction in the antitumor functionality of intILCs as they transition from the environment of a healthy liver to tumor-infested tissues.

### Downregulated ILC1 signature related to poor prognosis in liver cancer patients

Our data suggested that ILC1s were the most significantly reduced population in the HCC microenvironment, and expression of ILC1 signatures were decreased in all NK subsets. Thus, we predicted that the signature from ILC1 cluster would provide critical prognostic information in cancer patients. The 9 selected genes (*XCL1*, *CD160*, *CCL3*, *EOMES*, *IL2RB*, *CXCR6*, *XCL2*, *KLRB1*, *KLRC1*) were used to form the ILC1 signature. Analyses of cancer patient cohorts from TCGA (The Cancer Genome Atlas) showed that the low ILC1 signature expression group correlated with a significantly poorer prognosis in HCC (LIHC, liver hepatocellular carcinoma) (Figure 7A) and other types of cancer (LUAD, lung adenocarcinoma; SKCM, cutaneous melanoma; SARC, sarcoma) (Figure 7, B-D). These analyses supported the role of ILC1 signatures as an important indicator of clinical outcomes in HCC patients.

**Figure. 7.**
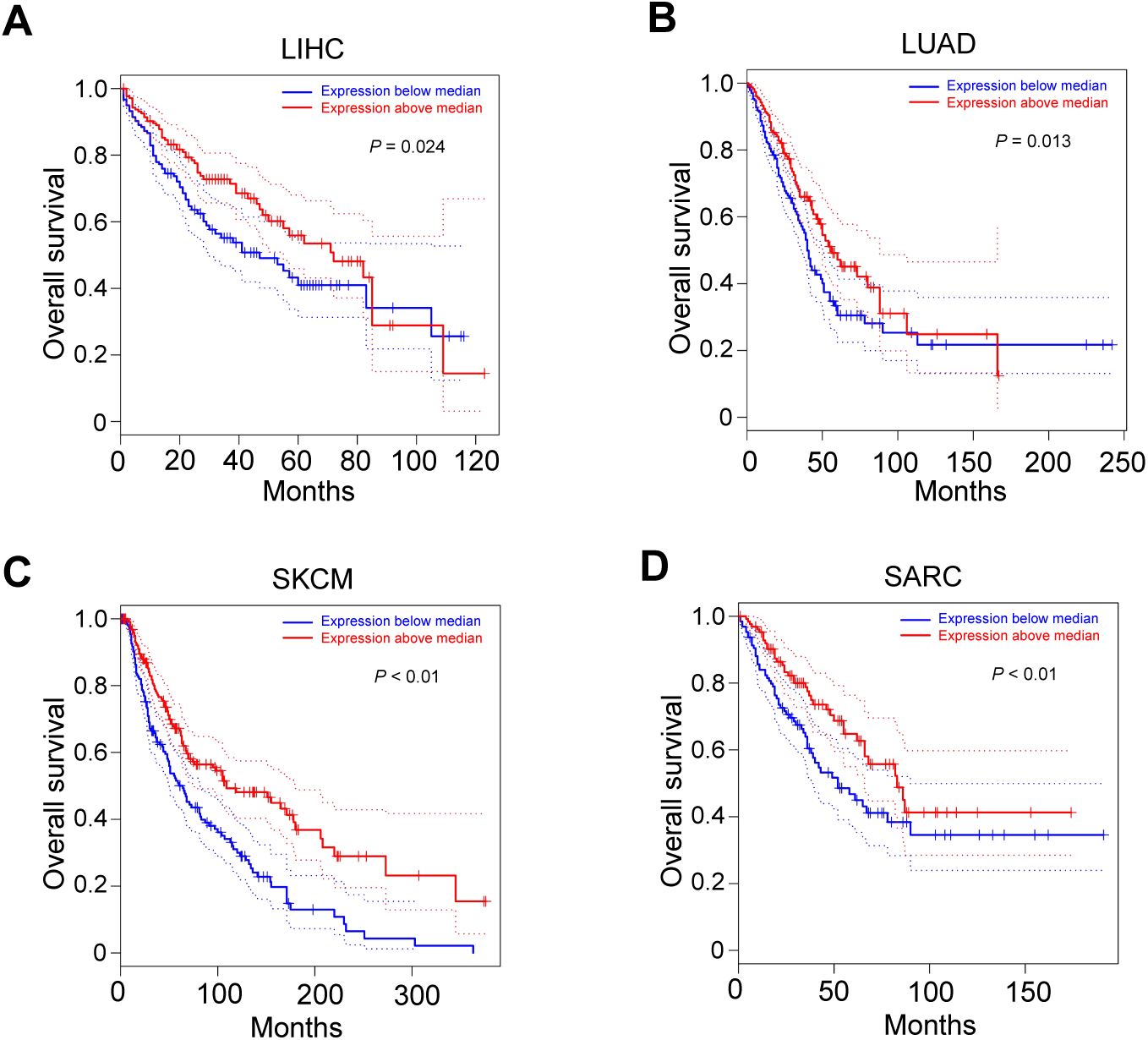
Prognostic role of ILC1 signature from single data. (A-D) Kaplan-Meier survival curves for overall survival from HCC **(A)** (LIHC, n=364); lung cancer **(B)** (LUAD, n=478); skin melanoma **(C)** (SKCM, n=458), and sarcoma **(D)** (SARC, n=262) samples from TCGA dataset, showing prognostic significance of ILC1 signature derived from scRNA-seq data.

## Discussion

Currently, identifying human liver ILC1 was a major challenge to distinguish human liver ILC1 from NK cells and to further study the individual function of ILC1. Varying from ILC1 in mouse liver, which has a well-defined definition, human liver ILC1 has an important problem that the identification criteria cannot be uniform, also making it difficult to study the characteristics of these cells. We identified the corresponding human liver ILC1(hILC1) according to mouse liver ILC1(mILC1) through integrated analysis of human and mouse liver single cell sequencing data, and verified the identity of this group of human liver ILC1 by DEGs and enrichment analysis, making a new attempt for the identification of human liver ILC1.

We successfully defined hILC1 and analyzed its differences from hNK cells, which have similar functional characteristics to mILC1. However, in terms of the expression of important cytokines Eomes, we found significant differences between hILC1 and mILC1: *EOMES* was low expressed in hNK, but overexpressed in hILC1; In mILC1 and mNK, the opposite is true. T-bet and Eomes are crucial transcription factors that regulate group L innate lymphoid cells, and were considered as two pivotal checkpoints in NK cell maturation[45]. In mice, both mNK cells and mILC1 require T-bet. However, *Eomes* deficiency specifically impacts the development of mNK cells, particularly the maturation of NK cells, whereas Eomes is dispensable for ILC1 development in mice[22]. This distinction between mNK cells and mILC1s, based on the dependency on Eomes, highlights a significant divergence in their maturation processes driven by transcription factors. Such insights have facilitated the generation of mouse models with mNK cell-specific deletions, leaving mILC1s unaffected[46]. Some studies found that NK cell development undergoes a transition from TRAIL^+^DX5^-^ to TRAIL^-^DX5^+^ stages. During this process, NK cells first require the expression of T-bet to acquire TRAIL^+^ expression. Following this phase, expression of EOMES is essential for the cells to achieve full functional maturity[45]. However, later studies found that in group L ILCs, especially in mouse liver, TRAIL^+^DX5^-^ ILC1s and TRAIL^+^DX5^+^ NK cells are actually two groups of relatively independent cells in development[21]. We used the mouse NK cells knockout *Eomes* by conditional knockout technique and the human NK cells knockout *EOMES* by CRISPR to investigate differences in transcriptome regulation of *Eomes(EOMES)* between mouse/human liver NK cells and ILC1s. In fact, the regulatory role of *EOMES* in human liver hNK and hILC1 cells may not be as important as observed in murine models.

We also investigated *Eomes* expression in NK cells and ILC1s in the liver of some other animals. T-bet^+^Eomes^+^ NK cells and T-bet^+^Eomes^-^ ILC1s in rat livers are thought to correspond to NK cells and ILC1s in mouse livers, but their expressions levels of CD49a and CD49b are different from those in mice: In rat ILC1, the proportion of CD49a^+^CD49b^+^ subgroup accounted for 75%, while in rat NK, the proportion of Cd49a^+^Cd49b^+^ subgroup accounted for about 40% and the proportion of CD49a^-^CD49b^-^ subgroup accounted for about 26%[47]. A population of NK cells in pig liver showed an Eomes^hi^T-bet^lo^ expression pattern distinct from mouse lrNK cells (ILC1) but highly similar to human lrNK counterpart. Furthermore, the porcine Eomes^hi^T-bet^lo^ liver NK cell population is able to produce IFN-γ upon IL-2/12/18 stimulation but lacks the ability to kill K562 or pseudorabies virus-infected target cells[48]. This low cytotoxicity is not like conventional NK cells, but more like ILC1. Therefore, the expression pattern and regulatory role of *Eomes (EOMES)* in NK cells and ILC1s may be different in different species.

NK cells and ILC1s, particularly liver-resident ILC1s, exhibit disparate developmental trajectories and originate from distinct progenitors[21]. Notwithstanding, transformation from NK cells to ILC1s has been documented in murine models of hepatic cancer[49]. In our investigations, while no discernible shift was noted in the proportion of NK cells or ILC1s within human hepatic tumors, significant alterations in the subpopulations of these cells were observed in the livers of patients with tumors. NK cells and ILC1s display intrinsic heterogeneity within healthy hepatic tissues, a characteristic that is conspicuously absent in both tumor and peritumoral tissues. In these environments, the persistence of a singular subpopulation is evident, which frequently lacks the canonical functions associated with NK cells or ILC1s, thereby favoring tumor progression through the suppression of immune surveillance. This observation suggests the potential for more intricate immunological interactions within the human liver. Taking advantage of the high-resolution capabilities of scRNA-seq, we have identified a novel subset of intermediate innate lymphoid cells (intILCs) within the liver. Unlike the near-total loss of polymorphism in NK cells and ILC1s within tumoral and adjacent non-tumoral tissues, two subgroups of intILCs are consistently present across healthy liver, tumors, and tumor-adjacent tissues. The functional implications of intILCs in normal hepatic physiology and their sustained presence in HCC warrant further exploration.

## Acknowledgments

This work was supported by National Key Research and Development Plan of China (2022YFF1202901); National Natural Science Foundation of China (82372801); The Zhejiang Provincial Natural Science Foundation of China (LY21H150002); Natural Science Foundation of Tianjin (23JCYBJC01560, 23JCYBJC01370, 21JCZDJC00430); and Science and Technology Planning Project of Tianjin Municipal Education Commission (2022YGYB14).

## Data availability statement

All relevant data are provided in the paper and supporting information file. All other data are available from the corresponding authors upon reasonable request.

## Author contribution

Yumo Zhang, Jitian He, and Youwei Wang designed the study, analyzed the data and wrote the manuscript. Zhouxin Yang and Youwei Wang interpreted results. The manuscript have been reviewed and approved by all authors.

