## supplemental Figures for "Dissecting the Heterogeneity and Tumor-Associated Dynamics of Human Liver Group I ILC via scRNA Sequencing Data"

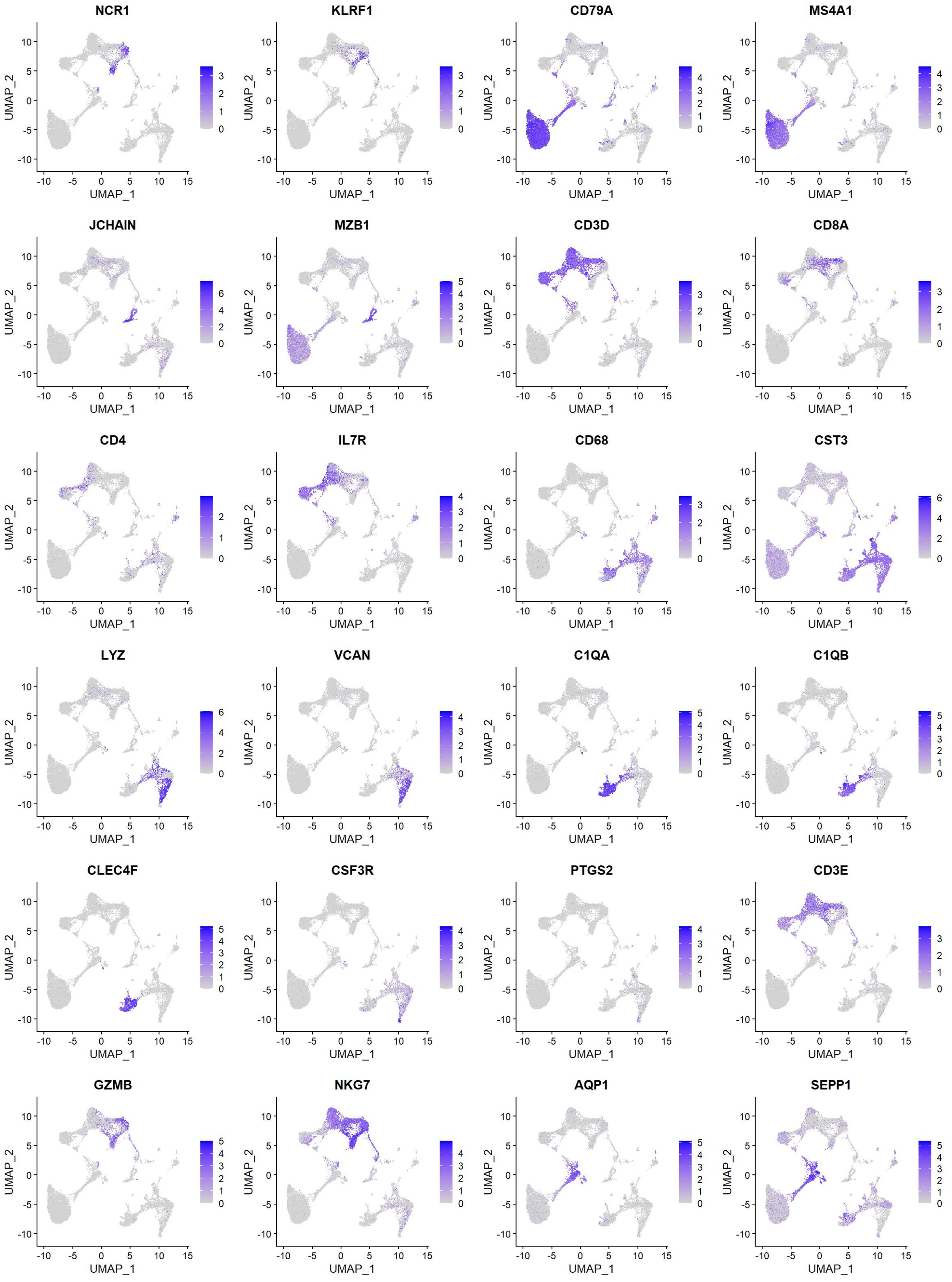


**Figure.S1 Representative marker genes of different cell clusters**
Feature plots showing the normalized expression of selected markers for all cell populations.

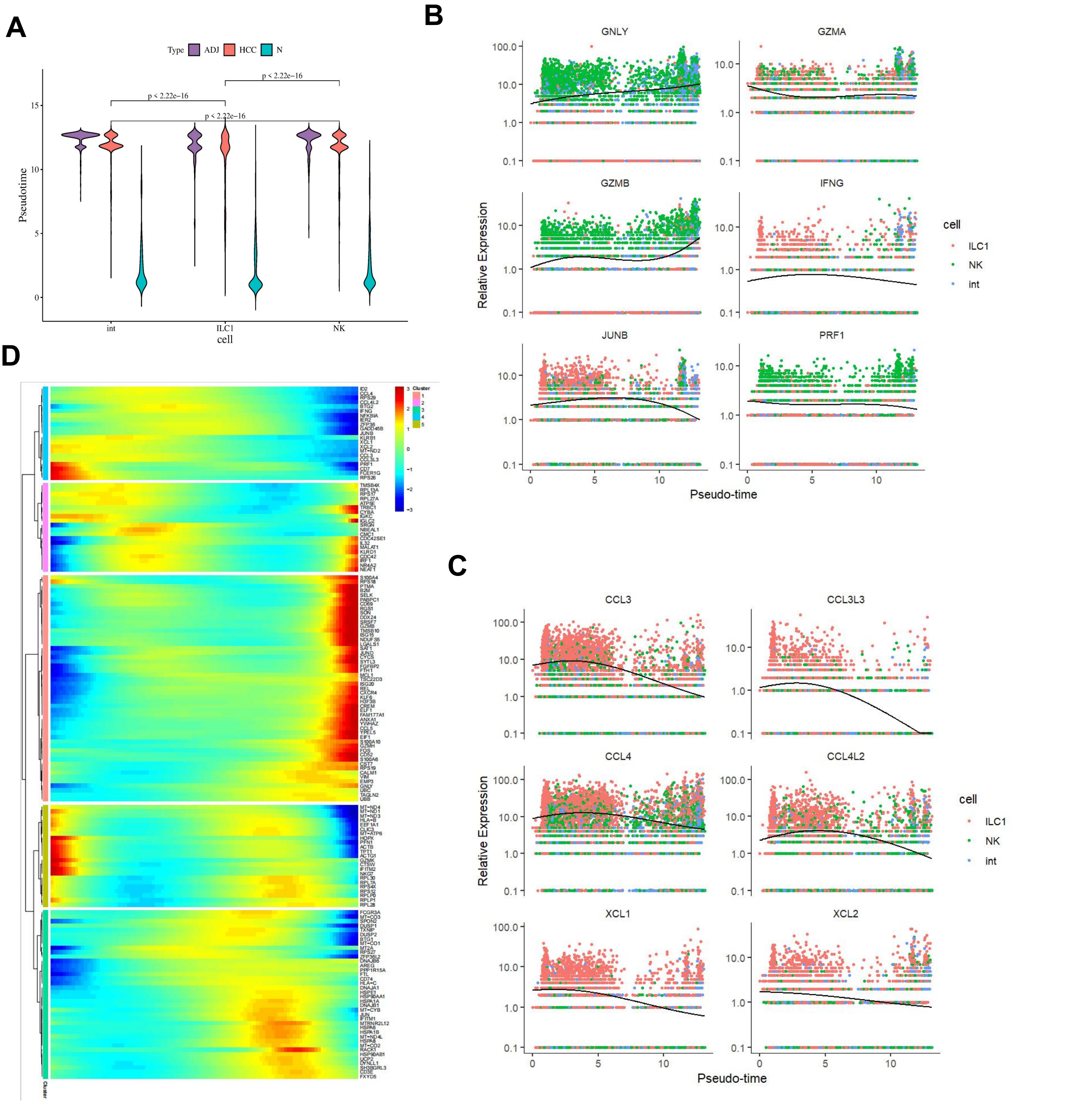


**Figure.S2 Pseudotime analysis of type 1 ILCs of HCC, adjacent tissue and healthy liver.**

**(A)** Violin plots showing the pseudotime value of NK cells, ILC1s, intILCs in 3 sample groups. **(B and C)** Two-dimensional plots showing the dynamic expression of some functional genes **(B)** and chemokine/ chemokine ligand genes **(C)** during the type 1 ILCS transitions along the pseudotime. **(D)** Heatmap showing all the cell-state transitions related genes which were divided into five clusters in Monocle2 analysis.

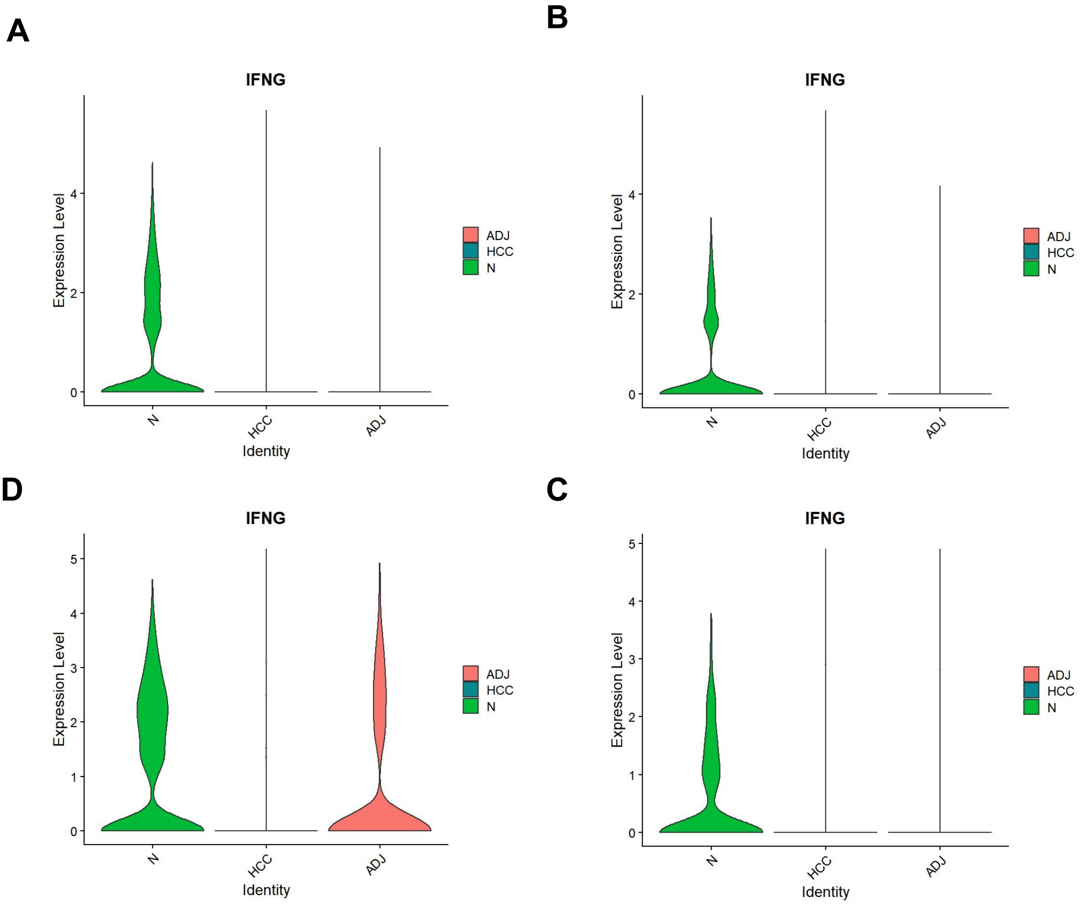


**Figure.S3 Violin plots showing the expression of *IFNG* in type 1 ILCs (A) , NK cells (B) , ILC1s (C) and intILCs (D) .**
